# A Mathematical Framework for Measuring and Tuning Tempo in Developmental Gene Regulatory Networks

**DOI:** 10.1101/2023.12.18.572181

**Authors:** Charlotte L Manser, Ruben Perez-Carrasco

## Abstract

Embryo development is a dynamic process governed by the regulation of timing and sequences of gene expression, which control the proper growth of the organism. While many genetic programs coordinating these sequences are common across species, the timescales of gene expression can vary significantly among different organisms. Currently, substantial experimental efforts are focused on identifying molecular mechanisms that control these temporal aspects. In contrast, the capacity of established mathematical models to incorporate tempo control while maintaining the same dynamical landscape remains less understood. This manuscript addresses this gap by developing a mathematical framework that links the functionality of developmental programs to the corresponding gene expression orbits (or landscapes). This unlocks the ability to find tempo differences as perturbations in the dynamical system that preserve its orbits. We demonstrate that this framework allows for the prediction of molecular mechanisms governing tempo, through both numerical and analytical methods. Our exploration includes two case studies: a generic network featuring coupled production and degradation, and the repressilator. In the latter, we illustrate how altering the dimerisation rates of transcription factors can decouple the tempo from the shape of the resulting orbits. The manuscript concludes by highlighting how the identification of orthogonal molecular mechanisms for tempo control can inform the design of circuits with specific orbits and tempos.

## 1 Introduction

Biological systems are intrinsically dynamic. Cells are constantly migrating, differentiating, dividing, and dying; collectively underpinning numerous biological processes. This dynamism is especially apparent during embryonic development, where the precise coordination of cell differentiation plays a pivotal role for an orchestrated formation of distinct tissues constituting the anatomical structure of the organism [1, 2, 3, 4, 5]. This coordination takes place across different chemo-mechanical scales and is locally encoded at the cellular level through genetic programs. Also known as gene regulatory networks, these programs confer the cell with the dynamical properties required for different cell differentiation processes [6, 7]. Notably, any inaccuracies in the timing of cellular dynamics can lead to critical errors that disrupt the healthy development of the embryo [1, 2]. Conversely, flexibility in the timing between tissues allows for the plasticity required for evolutionary traits to emerge. Given the crucial role of timing, a fundamental question arises in the field of developmental biology: How is the timing of cellular decisions precisely controlled?

One strategy to answer this question involves comparing homologous developmental processes across different species [8], particularly during the phylotypic stage when embryos closely resemble each other [9]. These processes usually preserve the same function across species, manifested as the same sequential pattern of activation of gene expression. Recent research has shown that these developmental programs, with evolutionary conserved cis-regulatory architectures, manifest different species-specific timings [10, 11, 12]. These different timings can vary by an order of magnitude across distinct species, and can be observed in processes like motor neuron differentiation [13], the segmentation clock [14, 15], and cortiogenesis [16]. During each one of these processes, the same genes are expressed in the same sequential manner, but at different speeds. Drawing an analogy from music, the species-specific execution speed of otherwise analogous genetic programs has been termed the *tempo* of the system [13, 17, 18, 19].

Furthermore, the species-specific tempo in these processes can be inherently cell-intrinsic, not requiring intercellular communication [5, 20, 21]. This observation has resulted in an extensive search for the intracellular mechanisms governing tempo, with recent candidates focusing on specific molecular steps of gene regulation such as protein stability, mRNA splicing, phosphorylation, or ubiquitination [13, 15, 22, 23, 16, 24]. Contributing to this complexity, all these different mechanisms can affect each other which, given the inherent non-linearity of gene regulation, impedes a straightforward mechanistic identification of tempo control.

The impact of non-linear feedback in gene regulation has been efficiently tackled in the past through mathematical modelling within the dynamical systems framework. This strategy offers a mechanistic insight into gene expression dynamics by establishing the rules governing reaction rates among biochemical species. The success of mathematical models resides in their ability to connect molecular mechanisms with a general understanding of the compatible dynamics [25, 26, 27, 28, 29]. Yet, the vast majority of the dynamical systems literature has been centered around steady-states, usually associated with available cell types, and very few papers focus on the transient dynamics leading to those states [30, 31]. This gap highlights the need for novel tools to comprehend the non-linear mechanisms governing tempo variations across species. In this manuscript, we devise a mathematical approach that connects the dynamic sequence of relative gene expression levels with the underlying biomolecular processes.

## 2 Matertials and Methods

### 2.1 Orbital equivalence: preserving the function of gene regulatory programs

Before attempting to compare quantitatively the tempo between two species, it is necessary to ask: can they actually be compared? Similarly, when we claim that the differentiation of two cells is the same except for their tempo, what do we mean by *the same*? In essence, we need to identify the differential characteristics of two temporal gene expression profiles that allow us to assert that they perform the same function but at a different tempo. Within the context of cell differentiation, the function of a genetic program is translated into the conserved sequence of relative expression of different genes, independently of the speed at which this sequence is travelled. Adopting the jargon of dynamical systems to describe this situation: two cells can have different gene expression trajectories, while still following the same orbits of gene expression. The orbits of the system can be visualised as the continuous sequence of cell states in the gene expression phase-space (see Fig. 1A and B).

**Figure 1:**
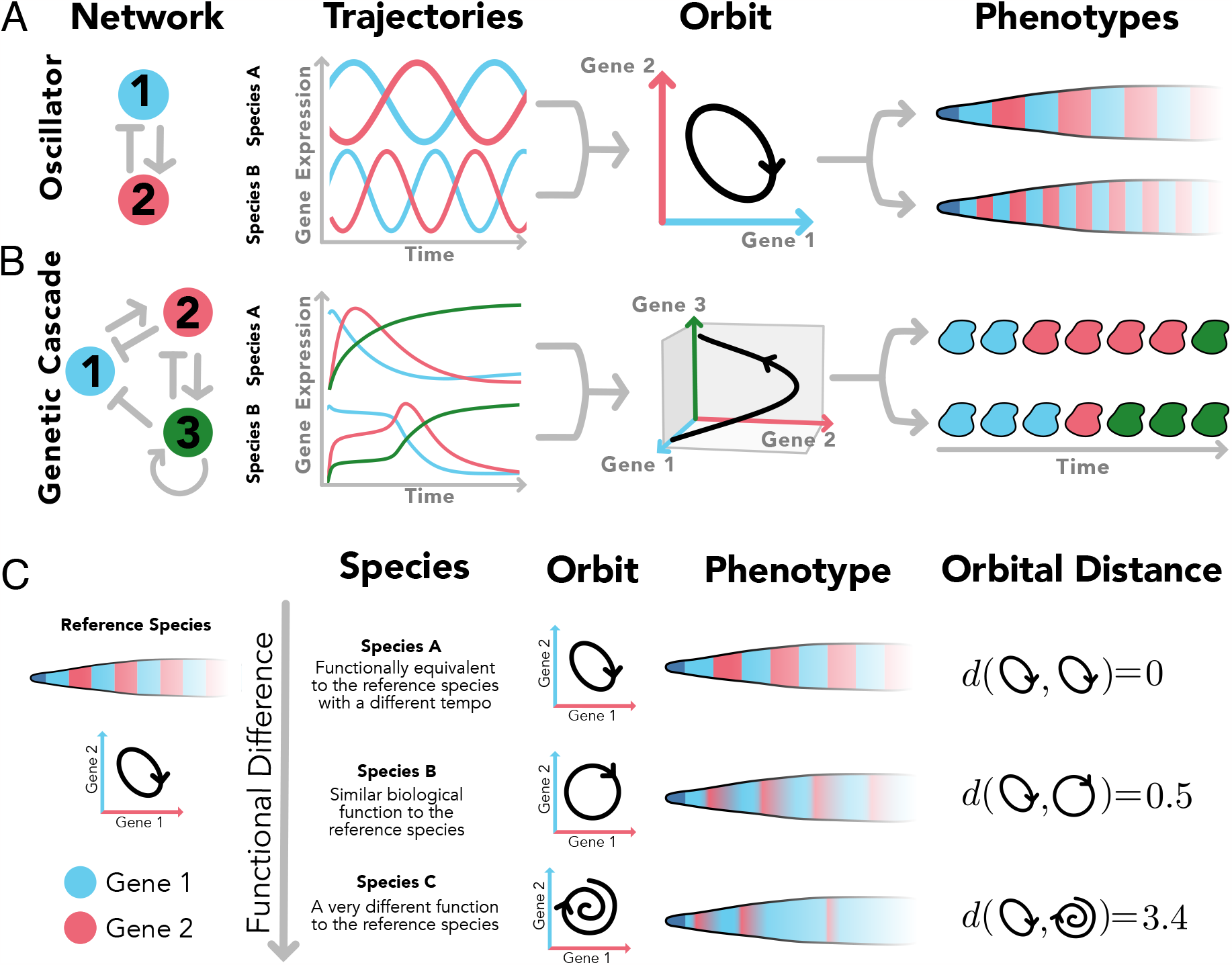
Orbital distance allows to compare the phenotypic function of two species. A-B) Two species (systems) that show the same relative sequences of gene expression in time will share the same orbit even though they might have different trajectories. This is translated in a different tempo while preserving the phenotypic function of the system. This is shown for an oscillator in charge of some spatial patterning (A) or more complex networks in charge of a cascade of gene activations (B). C) Distance between phenotypic functions for different species can be studied by comparing the orbits of the species defining an orbital distance *d*.

This identification of gene expression orbits with their core functional activity harmonizes with the proposed concept of dynamical modularity proposed by Jaeger and Monk [32]. We can employ this definition to compare how close the functions of two different dynamical systems are by quantifying the similarity of their gene expression orbits (see Fig. 1C). This strategy also reframes our exploration of tempo-controlling mechanisms: we can search for a suite of biochemical perturbations that sustain the integrity of a system’s orbit. Thus, the effect of these biochemical perturbations can exclusively change the tempo. In addition, given the ability of mathematical models to incorporate different levels of biochemical detail in gene expression, we can use this framework to compare how different molecular mechanisms (e.g. post-translational modifications or promoter occupancy) affect the orbits of expression of given genes.

Genetic networks can be modelled in a variety of ways, such as with discrete or stochastic descriptions [33]. For the purpose of this study, we will describe gene expression as a set of biochemical reactions that can be modelled as a system of ordinary differential equations [34, 35, 36]. In this description, the evolution of *N* different biochemical species in time 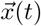 with 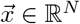 results by describing their rates of change,

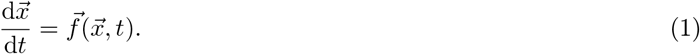

The functions 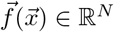 encapsulate the biochemical reactions describing the system. In general, to obtain the orbits of a genetic program, numerical integration of Eq. 1 is required. However, we can identify if two systems have the same orbits by directly inspecting their systems of equations. Specifically, when considering a second dynamical system given by 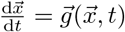 with 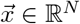, both systems share identical orbits if

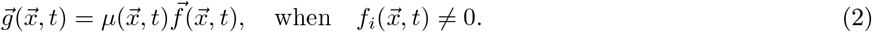

In other words, two systems have the same orbits when a scalar prefactor 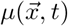 exists that scales the rates of change of the different biochemical species [37]. When this occurs it is said that both systems are *orbitally equivalent*. The prefactor 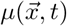 has a straightforward interpretation. It serves as the constant that scales the rate of change for each gene locally at any given specific gene expression state. This scaling relationship is shared among all the genes 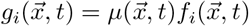 for genes *i* = 1,…, *N*. This preserves the overall direction of the rate vector 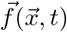 in the phase-space while keeping intact the resulting orbits (see Fig. 2A and B). In the context of dynamical landscapes, commonly used to understand differentiation trajectories [38, 39, 40, 41], the prefactor 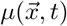 encodes all the possible changes that can be performed to the dynamical system that preserve intact the landscape.

**Figure 2:**
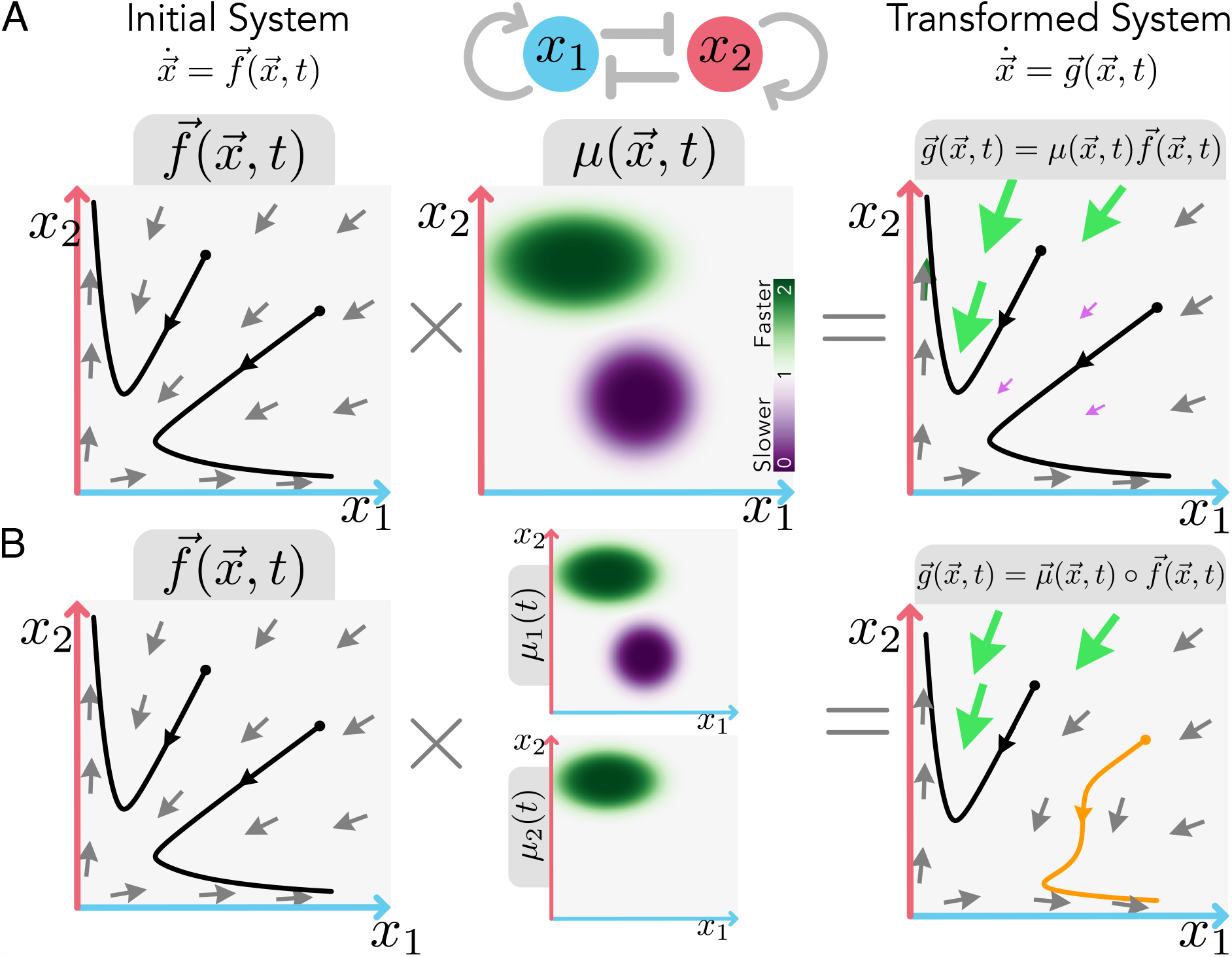
Orbital equivalence dissociates dynamical landscape and tempo between two dynamical systems. A) Schematic showing how a dynamical system 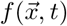 (left) for a given gene regulatory network (top) can be transformed into a different system that preserves the same orbits 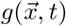 (right), by multiplying the initial system by a prefactor 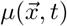 that preserves the dynamical landscape. Arrows show the flow of 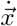. Black lines show two orbits for initial conditions given by the black. The prefactor might vary arbitrarily in space and time, giving rise to zones of the gene expression landscape that are faster (green arrows and zones), or slower (purple arrows and zones), while keeping the same orientation of the evolution of the system 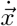 at any point 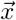. B) If the prefactor is not the same for every gene (so that it is represented as a vector 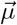 rather than a scalar *μ*), the resulting orbits may not be preserved. In this scenario, the system 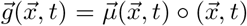 results from the element-wise multiplication (also known as Hadamard product) for each component 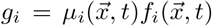. Prefactor heterogeneity among genes only impacts orbits in gene expression zones where the prefactors differ (depicted by the orange orbit). Parameters and functions used can be found in the SI.

Moreover, 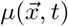 quantifies the relative tempo of a particular process given by its harmonic mean along the orbit of that process, 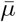, written as the ratio of line integrals (see SI for a detailed derivation),

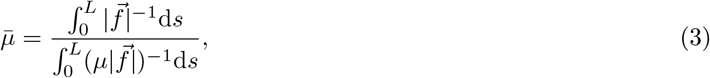

where *L* is the total length of the orbit in the state space. Note that orbital equivalence leaves plenty of freedom to the form of the prefactor 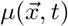, which can change arbitrarily in the gene expression space (see Fig. 2A). Hence, under the lens of dynamical systems, it becomes apparent that tempo changes are not limited to global constant organism-specific tempos, but more generally to changes in the local speed (dependent on the gene expression state 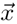) at which the same gene dynamical landscape is traversed.

### 2.2 Orbital distance: comparing the function of gene regulatory programs

In general, it is not anticipated that two organisms will exhibit precisely identical orbits, as slight variations in orbits will maintain the overall function of the cell differentiation program (see Fig. 1C). Similarly, changes in biochemical rates are not expected to result in exact orbital equivalence (Eq. 2). Taken together, this highlights the requirement to define an orbital distance that measures the divergence between orbits in different systems. Along this manuscript, we will evaluate the distance of two orbits using the Fréchet distance, which captures the maximum deviation between two orbits, becoming zero when two systems are orbitally equivalent. Several alternative definitions of orbital distance are possible [42, 43], each able to capture different aspects of the similarity between two different gene expression sequences (see SI and Fig. S.1 for more examples of possible measures). Ultimately, the choice of orbital distance needs to address how similar the function of two systems is. This is not only required to compare two mathematical models but also applies to the comparison of two experimental datasets of gene expression when it is claimed that two species only differ in their tempo (regardless of whether the underlying GRN is known).

In addition, when investigating the role of different molecular mechanisms using the dynamical systems modelling framework, we can also attempt to define an analytical orbital distance using the definition of orbital equivalence (recalling Eq. (2)). This can be done by searching for parameter modifications that can be written as a multiplicative prefactor in the dynamical system. In general, it is not expected that arbitrary perturbations will entail exact orbital equivalence. Instead, changes in biochemical parameters that preserve the function of the system will involve a gene-dependent prefactor (i.e. a vector 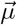 that is no longer the same scalar for each single gene) that is comparable across different genes. The new system follows the element-wise (or Hadamard) product 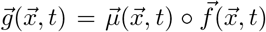, where 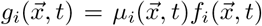. When the components of the prefactor are different from each other, the systems will not be orbitally equivalent (see Fig. 2B). Hence, an orbital distance can be defined by quantifying the heterogeneity of the components of the prefactor vector 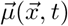. This can be achieved through measures like the weighted variance,

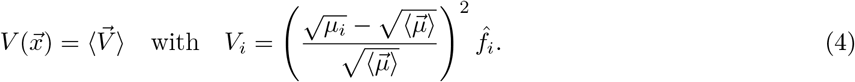

Where 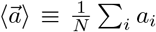 is the average taken over the components of a vector 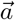. The weights 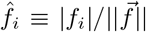 correspond to the normalised velocity vector’s components (Eq. 1). These weights ensure that heterogeneity is accounted for in the molecular species that shape the system’s orbit at certain points in the state space. For instance, a prefactor component *μ*_*j*_ different from the system’s average ⟨*μ*_*i*_⟩ will not compromise orbital equivalence within regions of the state space where that gene expression is not changing *f*_*j*_≃ 0. Finally, we want to evaluate the heterogeneity 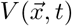along the system’s orbit. To compare with the results from the Fréchet distance, we can define an orbital prefactor heterogeneity distance as the maximum heterogeneity along the orbit,

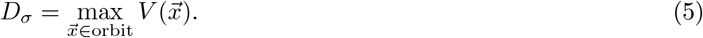

For two identical orbits *D*_*σ*_ = 0, while *D*_*σ*_ grows as the prefactors become more dissimilar. Together with our measure of tempo (Eq. 3), we can interrogate how different mechanisms influence these factors by searching for mechanisms where *D*_*σ*_ is minimised, while the tempo is allowed to vary.

## 3 Results

### 3.1 Scaling production and degradation

One immediate result from the orbital equivalence framework is that identical global rescaling of protein production and degradation rates of the transcription factors of a given gene regulatory network will preserve the orbit while controlling tempo. Interestingly, this rescaling does not have to be constant in time but can change arbitrarily in time (see Fig. 3). This provides a mechanism to preserve the orbit of gene expression independently of changes along the cell cycle or external perturbations as long as production and degradation are rescaled in the same way. This result applies to any given regulatory network independently of the complexity of its topology (see Fig. 3), suggesting a robust mechanism to tune tempo of a regulatory system, allowing for evolutionary strategies.

**Figure 3:**
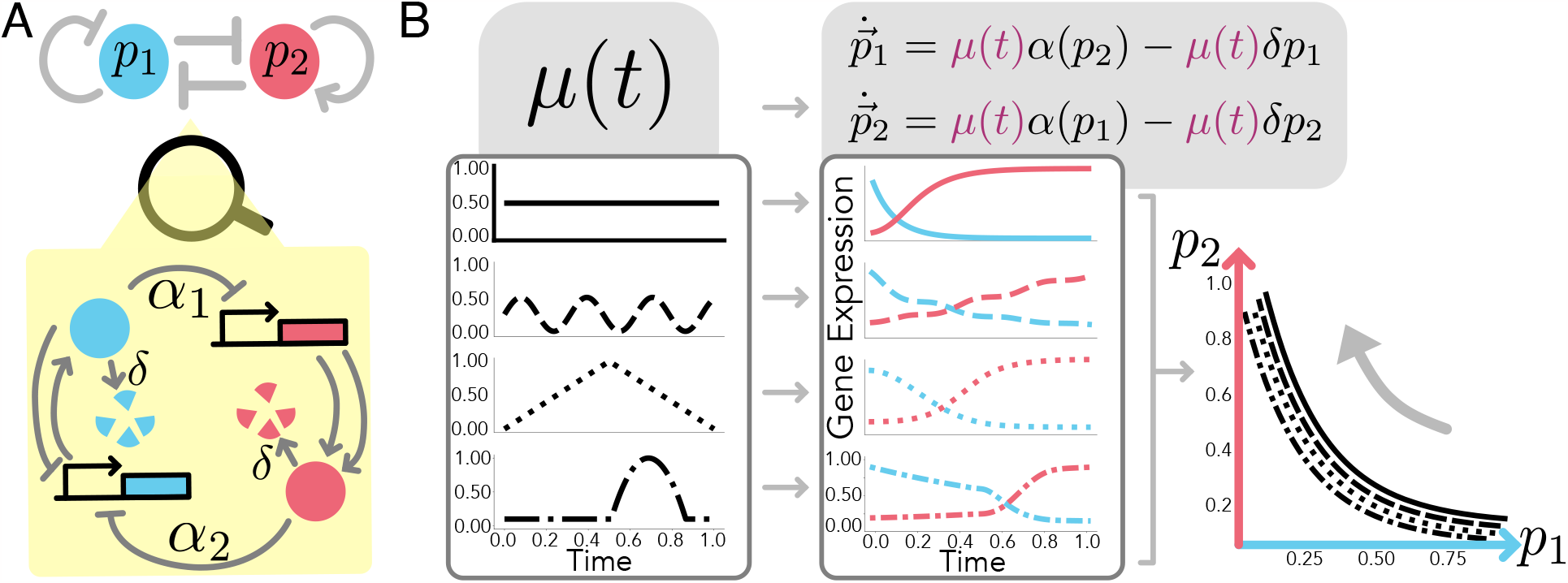
Scaling of production and degradation can control tempo. A) A gene regulatory network which produces a simple 2-gene genetic cascade (a switch from high expression of *p*1 to high expression of *p*2), together with a molecular schematic of the mechanisms involved. B) Using various arbitrary prefactors *μ*(*t*) (left) which scale the protein production and degradation, we can affect the trajectory of the GRN (centre) while keeping the same orbit (right). This effectively changes the switching tempo while keeping the relative expression of the genes along the switch. Parameters and functions used can be found in the SI.

### 3.2 Case study: The Repressilator

In general, arbitrary perturbations other than global scalings of production and degradation will not necessarily result in orbital equivalence. To showcase the applicability of the orbital equivalence framework in these scenarios, we will implement it within a well-known gene regulatory network, the repressilator (see Fig. 4A). Despite not corresponding with particular developmental biology example, it is a paradigmatic circuit capable of gene expression oscillations that offer the flexibility to incorporate different degrees of biochemical complexity [44, 45, 46] and different tempos [47]. In addition, sustained oscillations result in closed orbits in the state-space (see Fig. 4B and C), allowing for a clearer visualization of the concept of orbital distance.

**Figure 4:**
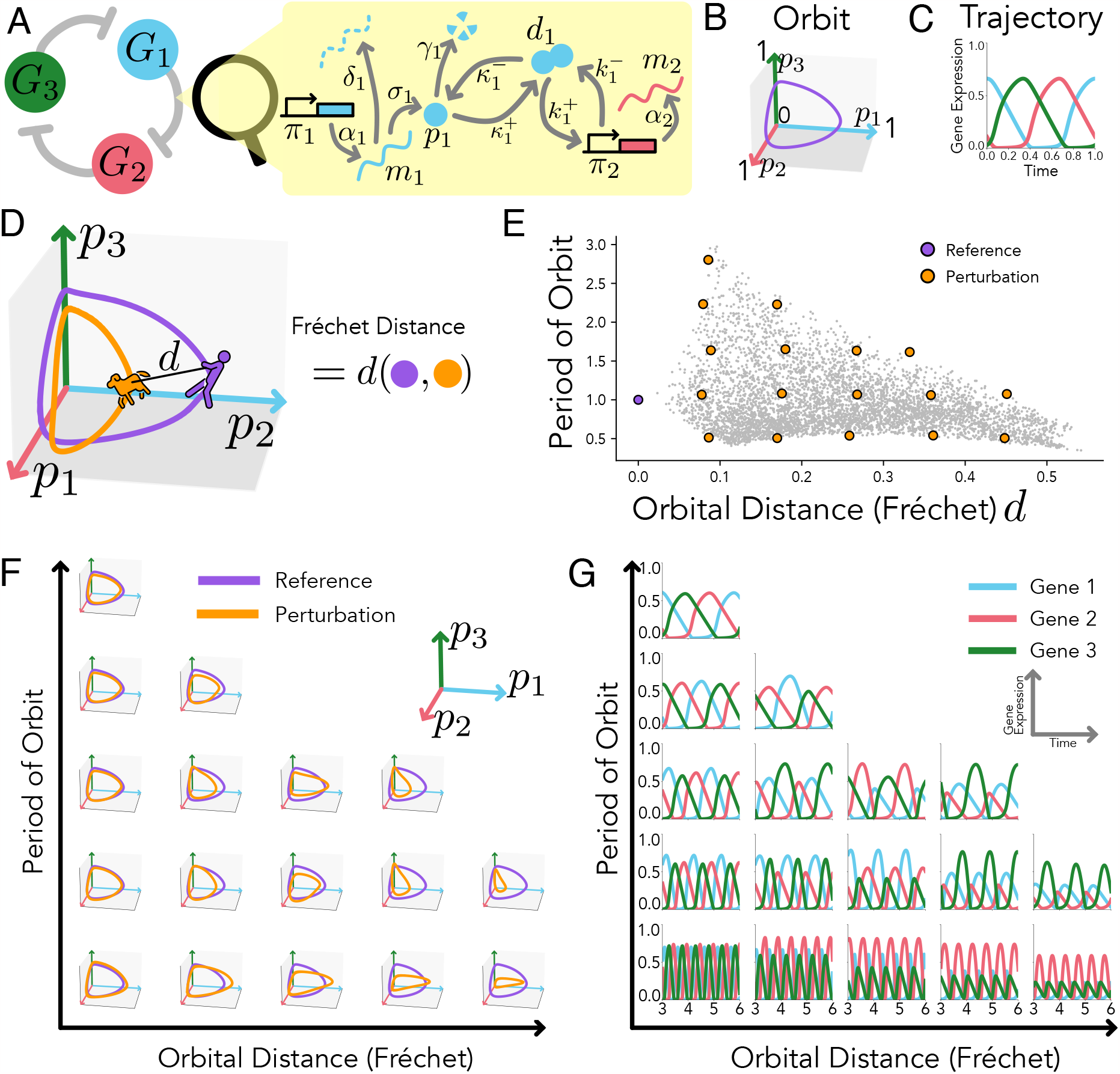
Molecular perturbations can affect orbit and the tempo. A) A schematic of the repressilator in which three genes repress each other in turn (left) indicating the intermediate biochemical steps considered. B-C) The repressilator can give rise to stable oscillations of gene expression that result in closed orbits (B) and periodic trajectories (C). D) The orbital distance between two different systems is measured with the Fréchet distance, a measure of similarity between two curves. The Fréchet distance can be intuitively understood as the shortest lead distance *d* needed for a person walking a dog, where the person walks along one curve, and the dog walks along the other, allowing both to move at independent speeds. E) Resulting period and orbital distance of 5000 randomly selected perturbed systems (grey dots) with respect to a reference system (purple dot). For each parameter set, each rate 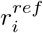 was perturbed with a uniform fold change returning new rates 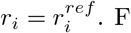) Orbits for selected perturbed parameter sets (corresponding with orange circles in E), confirm that some parameter sets can keep the orbit shape while changing tempo (period of the orbit), while others change the orbit of the system while keeping the same period of oscillation. G) Trajectories for the same selected parameter sets showing the perturbation effect in the temporal evolution of gene expression.

To explore the possibility of different post-transcriptional mechanisms we employ the molecular description presented by Bennett et al [48], containing an explicit dimerization and promoter occupancy (see Fig. 4A). In particular, each one of the 3 genes composing the circuit involves 4 distinct variables: mRNA (*m*_*i*_), protein (*p*_*i*_), protein dimer (*d*_*i*_), and promoter occupancy with two possible states with a probability of being active (*π*_*i*_). The evolution of these 12 variables is given by:

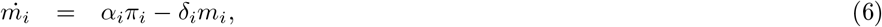

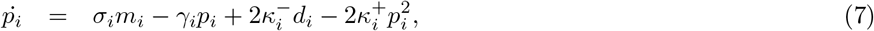

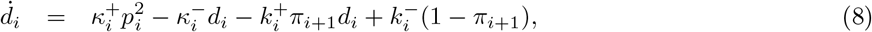

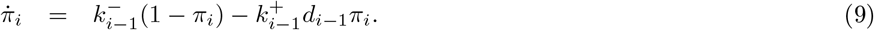

Where *i* = (1, 2, 3) is a cyclic index running through the three genes. This description contains 8 reactions for each gene, yielding a total of 24 different biochemical rates (see Fig. 4a): transcription (*α*_*i*_), translation (*σ*_*i*_), degradation of mRNA (*δ*_*i*_), degradation of protein (*γ*_*i*_), reversible dimerization 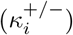, and reversible promoter binding of a dimer, which silences the transcription of the succeeding gene 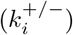.

This high-dimensional system can be reduced by leveraging the difference in timescales of the dynamics of the different biochemical species. Employing quasy-steady-state (QSSA) approximation methods [48, 49, 50, 51], we can simplify the system of equations by assuming that the change in time of total protein amount in any of its configurations (monomer, free dimer, or bound dimer) changes at a slower timescale than the relative differences between any of these possible configurations. Under this assumption we can reduce the system to only two sets of equations for each gene (see SI for more details):

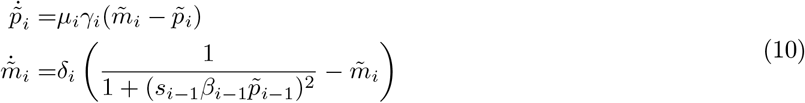

with,

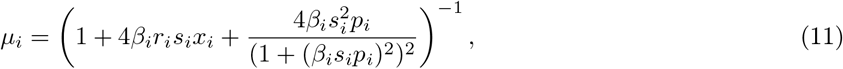

where we have introduced three new parameters that control the system dynamics 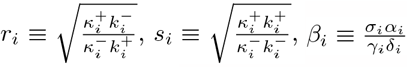. In particular *r*_*i*_ and *s*_*i*_ control the directionality and relative magnitude of the reversible intermediate reactions that connect the monomeric protein and the bound promoter. Since we are interested in the overall shape of the orbit regardless of the absolute number of molecules, without any loss of generality we have also rewritten the dynamics in terms of the non-dimensionalized mRNA and protein expression 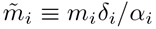 and 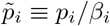. Finally, we can assume that mRNA dynamics reach equilibrium fast with respect to the value of 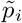. Applying QSSA to mRNA we can simplify the system further (see SI).

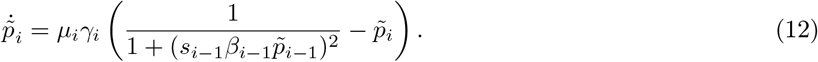

It is immediate to see that the effect of the intermediate protein mechanisms is summarized as a prefactor *μ*_*i*_. Comparing this result with Eq. (2), we can identify *μ*_*i*_ as the prefactor in charge of orbital equivalence. In particular, the only biochemical parameters that appear exclusively in the prefactor (see Eq. 11 and 12) are *r*_*i*_, controlling the reaction flux of free dimer, suggesting a route to control tempo. Since the three genes have different prefactors *μ*_*i*_(*p*_*i*_), that depend on the parameters and gene expression, we need to employ the different tools of orbital distance to assess the limits and characteristics of tempo control.

To explore the dynamical properties available through changes in the directionality of reactions towards the dimer state *r*_*i*_ we undertook a parameter search for different perturbations of the parameters *r*_*i*_ with respect to an arbitrarily chosen reference system. Analysis of the resulting oscillatory systems shows high variability in the possible oscillatory behaviours (Fig. 4D, E). The resulting orbits not only exhibit a large variability of shapes (Fig. 4F) but there is also variability in the periods of the orbits (Fig. 4G). Most interestingly, there are many cases in which the period of the orbit can be perturbed without changing the orbit of the system. Similarly, there are cases in which different orbits are obtained while preserving the period of the oscillations, suggesting that there are orthogonal molecular strategies to change orbit shape and tempo.

### 3.3 Identifying Molecular Mechanisms of Orbital Distance and Tempo

Observing the variability of behaviours accessible by changes in *r*_*i*_, we investigated whether we could predict the orbital distance and resulting period of the perturbed systems obtained in our parameter search by analytical inspection of the prefactors *μ*_*i*_ (Eq. 11). Using our analytical definition of prefactor heterogeneity *D*_*σ*_ (as outlined in Eq. 4 and Eq. 5) and comparing it with the orbital distance calculated by numerical integrations of the perturbed systems, we observed a linear relationship between the two (Fig. 5A). This relationship reveals an analytical means to predict changes in the orbit’s shape induced by a perturbation, without needing to integrate the perturbed ordinary differential equations.

**Figure 5:**
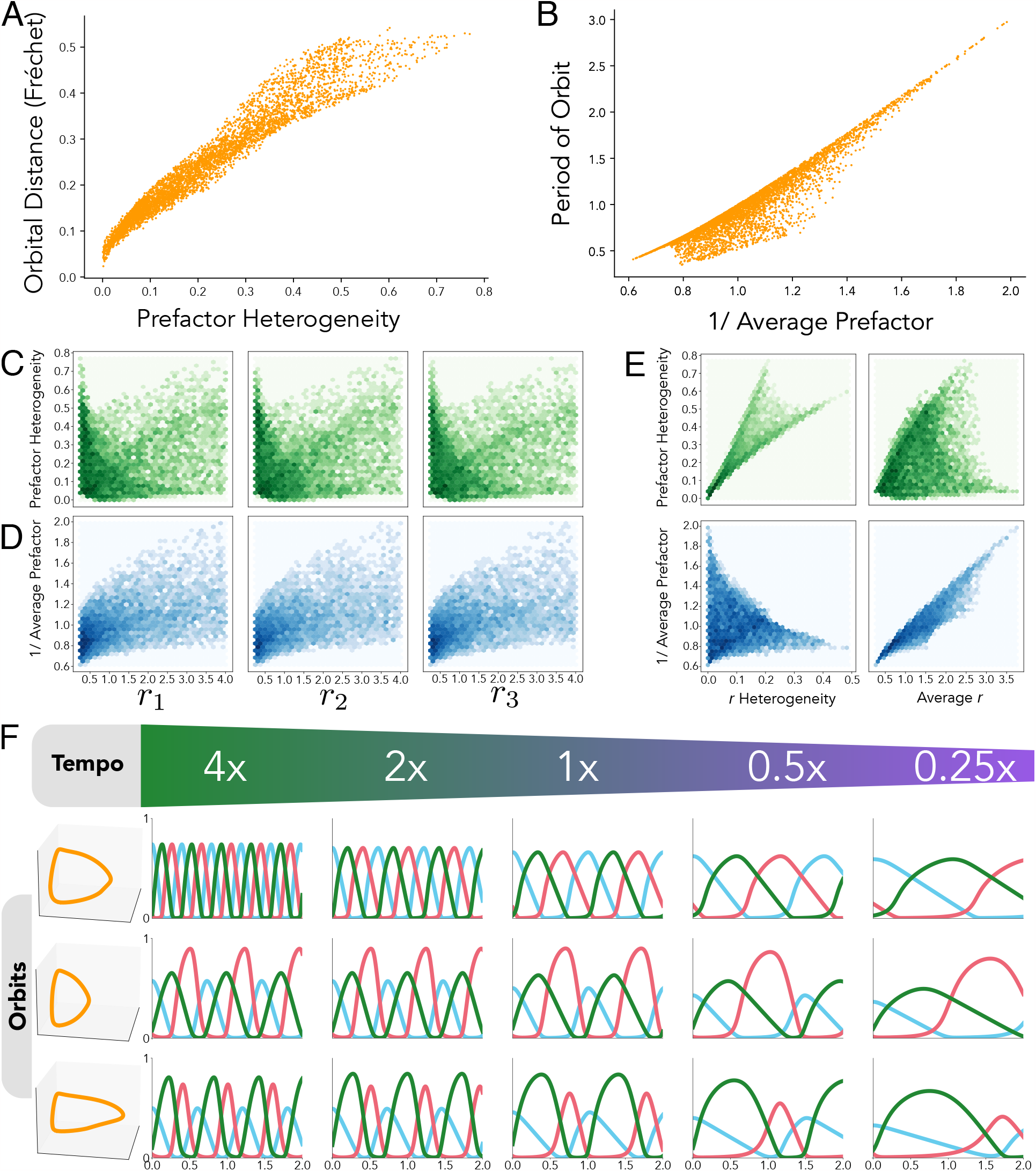
Identification of tempo mechanism reveal molecular strategies of tempo control. A) Prefactor heterogeneity (Eq. 4) for the sampled points in Fig. 4 serves as a good predictor for orbital distance (calculated as the Fréchet distance to the reference orbit), allowing us to predict if a parameter set will preserve the orbit without numerical integration. B) Fold change of the average prefactor with respect to the reference system calculated as 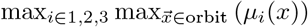, gives a good prediction of the fold change in the orbit’s period. C, D) Individual values of *r*_*i*_ are not enough to predict the prefactor heterogeneity (green), or the average prefactor (blue). E) The aggregate measurements *r* heterogeneity and mean(*r*_*i*_) are good predictors of the prefactor heterogeneity and the average prefactor respectively. F) For 3 different chosen orbits (rows), 5 different molecular perturbations (columns) are designed that result in a system that preserves the orbit while changing the tempo. The parameter sets were designed by changing the mean value of *r*_*i*_ while keeping their heterogeneity (see SI).

Similarly, we investigated whether the period of oscillations in the perturbed system could be predicted by analyzing only the prefactor of the new system. In this analysis, we found that the average prefactor serves as a reliable predictor of oscillation periods (Fig. 5B). This finding offers a method for forecasting the resulting oscillatory period without the numerical necessity of integrating the ordinary differential equations of the perturbed system.

While *μ*_*i*_ are good indicators of the effect of a perturbation in the system, they come with two main disadvantages. On one hand, the prefactor needs to be evaluated along the orbit of the reference system (Eq. 5), on the other hand, the prefactor is not immediately linked with the molecular mechanism in charge of orbital distance and tempo. Hence, we investigated if we can identify molecular mechanisms of tempo and orbit shape control with definitions that link directly to the combined rates towards dimerization *r*_*i*_. Initial inspection of the role that individual rates have on the prefactor reveals that, while there is some correlation between *r*_*i*_ and the resulting prefactor heterogeneity and average prefactor, there is not a strong predictive relationship (Fig. 5C, D). These results suggest that the effect that changes of prefactor have on the orbit originate from a combination of perturbations in the ensemble of rates included in *r*_*i*_. In particular, we identified that the average of the composed dimerization rates completely predicts the average prefactors, and consequently the period of the resulting orbit (Fig. 5E and Fig. S.2B). On the other hand, the heterogeneity of *r*_*i*_ allows us to predict the prefactor heterogeneity, resulting in a predictor of the orbital distance (Fig. 5E and SI Fig. S.2C). Unlike the orbital distance, or prefactor analysis (e.g. *D*_*σ*_), calculating *r*_*i*_ heterogeneity and average does not require any knowledge of the orbit of the reference system. Instead, it provides direct molecular mechanistic insight into the effects of post-translational rates on trajectories.

These results provide two orthogonal molecular mechanisms to control orbit shape and tempo. Hence, this allows us to use these mechanisms to design sets of parameters that will preserve the orbit of a system while exclusively changing the tempo of the system, independently of the reference system. We can achieve this by tuning the average *r*_*i*_ while fixing the heterogeneity in the prefactor. Implementing this strategy, we successfully gained the capability to fine-tune the tempo for orbits of various shapes (Fig. 5F), resulting in achievable speed differences spanning a full order of magnitude.

## 4 Discussion

In this paper, we have developed a mathematical framework using dynamical systems which can be used for investigating molecular mechanisms in charge of inter-species differences in developmental tempo. This framework is based on the notion of orbital distance, assigning a dynamical function to a system based on a defined progression of relative gene expression states. We have shown how we can explore this idea in two parallel ways: on one hand comparing orbits directly from gene expression data, on the other hand by providing an analytical toolkit connecting ordinary differential equation representations with developmental tempo.

The versatility of the techniques introduced means that they can be applied to many different genetic programs in development. The similarity measures based on gene expression data can be immediately used to compare function between organisms regardless of the underlying gene regulatory network. While this paper employs the Fréchet distance for this comparison, many other possible distances are possible (see SI), reflecting different definitions of function in embryogenesis.

The mathematical framework becomes very useful when contrasted with more traditional representations of the differentiation process as trajectories in a Waddingtonian landscape. Recent advances in this field have focused on reconstructing these landscapes from experimental data, usually fixing the location and change of steady states and separatrices of the landscape [38, 41]. Here we show how there are infinite available transformations of the dynamical system that preserve the same identical landscape. This transformation can be achieved through a prefactor of the dynamics (see Fig. 2A). Overall, this reveals two compatible formalisms that separate the geometry and the tempo of a dynamical system’s landscape. This not only allows us to understand the underlying mechanisms of tempo control but also informs how we interpret and find landscape representations of a particular differentiation process.

One of the main results of the manuscript is showcasing that tempo can be controlled through intermediate reaction steps during the regulatory process. Previous experimental studies have shown that inter-species developmental tempo depends heavily on specific biochemical parameters [15, 13, 11, 52]. Thus, the tools developed could be employed to investigate such systems to understand the impact and limits that different parameter perturbations will have on the resulting trajectories of gene expression. In order to do so, quasi-steady-state approximations need to be developed that can capture the impact that metabolic and other biochemical processes have on the overall change of gene expression in the state-space. In our case we identified dimerization rates as a pathway to control tempo, nevertheless we anticipate that other intermediate steps may have similar temporal implications.

All in all, this framework not only shows a route to quantitatively connect molecular mechanisms with tempo in development, but it can also be extended to other areas. For instance, in an evolutionary context, the same tools can be used to identify possible molecular routes to change tempo while preserving the function of a system. Hence, identifying evolutionary strategies that showcase the molecular requirements at different tempos. Similarly, these strategies can be employed directly in synthetic circuit design, with the aim of building circuits that allow for tempo control of specific functions, or even controlling the tempo of in vitro cell differentiation.

## Acknowledgements

We thank Dr. Smitha Maretvadakethope for her feedback on the manuscript. Our acknowledgements also extend to Prof. Nick Jones and Prof. Robert Endres for their insightful discussions about the project. C.M. gratefully acknowledges the financial support from the Department of Life Sciences at Imperial College London. R.P-C acknowledges funding from the Leverhulme Trust project grant RPG-2023-085. R.P-C also extends gratitude for the discussions at the KITP, which were supported in part by the National Science Foundation under Grants No. NSF PHY-1748958 and PHY-2309135.

