## Supplementary Information for "A Mathematical Framework for Measuring and Tuning Tempo in Developmental Gene Regulatory Networks"

### Tempo harmonic mean derivation

Given a dynamical system $\frac{d\vec{x}}{dt} = \vec{f}(\vec{x})$, the time $T_f$ it takes to travel a certain orbit $\mathcal{C}$ of arclength $L$, follows

$$T_f = \int_0^{T_f} \mathrm{d}t = \int_{\mathcal{C}} \left|\frac{\mathrm{d}t}{\mathrm{d}s}\right| \mathrm{d}s = \int_0^L |\vec{f}|^{-1} \mathrm{d}s. \tag{S.1}$$

A second system orbitally equivalent to the first $\frac{d\vec{x}}{dt} = \vec{g}(\vec{x}, t) = \mu(\vec{x})\vec{f}(\vec{x})$ will take a time $T_g$ following the same orbit $\mathcal{C}$

$$T_g = \int_0^{T_f} \mathrm{d}t = \int_0^L |\vec{g}|^{-1} \mathrm{d}s = \int_0^L |\mu\vec{f}|^{-1} \mathrm{d}s. \tag{S.2}$$

The relative tempo $\bar{\mu}$ between two orbits is the ratio of these two times

$$\bar{\mu} = \frac{T_f}{T_g} = \frac{\int_0^L |\vec{f}|^{-1} \mathrm{d}s}{\int_0^L (\mu|\vec{f}|)^{-1} \mathrm{d}s}. \tag{S.3}$$

### Alternative definitions of orbital distance

In this manuscript, we have employed the Fréchet distance to measure the discrepancy between two orbits. However, the literature offers a variety of alternative metrics for comparing curves. Some of these include Dynamic Time Warping (DTW), the Hausdorff Distance (HD), and the Distance Difference Matrix (DFD) [1] (Fig. S.1 A). These measures typically involve traversing both curves simultaneously and computing a distance metric during this process. For example, a common way of differentiating between curve distance methods is by classifying them as either bottleneck or aggregate methods. A bottleneck method is determined by finding the maximum point of difference between two curves. Meanwhile, an aggregate method calculates the distance as an average value along the entire length of both curves.

Another important consideration in selecting a distance metric is whether the comparison should be conducted by navigating both curves in a fixed sequential order (order dependent) or whether backtracking is permitted (order independent). This means that, when evaluating various orbits for their proximity to a reference orbit and arranging them in order, distinct curve distance measures may yield different rankings for the orbits (see Fig. S.1 C). Nevertheless, despite these differences, we observe similar results when using Fréchet, Hausdorff and DTW to compare against analytical measures of orbital distance through the prefactor (Fig. S.1 B).

### 2D Gene Regulation Equations

The system used to generate the bistable switch landscape of Fig. 2 in the main text is the following:

$$\begin{aligned} \dot{x} &= \mu(x,y)\left(\frac{1}{1+0.118\,(1+33y)^2} - x\right) \\ \dot{y} &= \mu(x,y)\left(\frac{1}{1+1.75\times 10^{-4}\,(1+1000x)^2} - y\right). \end{aligned} \tag{S.4}$$

The scalar prefactor $\mu(\vec{x})$ for Fig. 2A is based on the two ellipses

$$\begin{aligned} c_1 &: 0.25 = e_1(x,y) = 2(x-0.3)^2 + (y-0.75)^2 \quad \text{and} \\ c_2 &: 0.3 = e_2(x,y) = (x-0.5)^2 + (y-0.3)^2, \end{aligned} \tag{S.5}$$

where $c_1$ determines the zone with increased tempo (green area in Fig. 2), and $c_2$ the area where dynamics slow down (purple area in Fig. 2). Inside ellipse $c_1$ the tempo is increased following the prefactor,

$$\mu(x,y) = 1 + \exp\left(-10^{-4}(e_1(x,y))^2\right), \tag{S.6}$$

while inside ellipse $c_2$ the tempo is decreased following the prefactor

$$\mu(x,y) = 1 - \exp\left(-1.25 \times 10^{-3}(e_2(x,y))^2\right). \quad \text{(S.7)}$$

The prefactor is kept intact ($\mu(x,y) = 1$) outside of both ellipses. For the vector prefactor $\vec{\mu}(x,y) = (\mu_1(t), \mu_2(t))$ in Fig. 2B, $\mu_1(x,y)$ is the same as the scalar $\mu$ above. However, $\mu_2(t)$ does not include the slow-down zone $c_2$.

The incoherent feedback loop network used in Fig. 3 in the main text is the following:

$$\begin{aligned}
\dot{p}_1 &= \mu_n(t)\left(\left(\frac{1}{1+10{p_2}^2}\right)\left(\frac{1}{1+10{p_1}^2}\right) - 10p_1\right) \\
\dot{p}_2 &= \mu_n(t)\left(\frac{10+{p_2}^2}{1+6{p_1}^2} - 10p_2\right),
\end{aligned} \quad \text{(S.8)}$$

The prefactors $\mu_n(t)$ given by the profiles in Fig. 3 are the following:

$$\begin{aligned}
\mu_1(t) &= 1 \\
\mu_2(t) &= \frac{1}{4\sin(20t) + 1.1} \\
\mu_3(t) &= \min(2t, 2-2t) \\
\mu_4(t) &= \begin{cases} 0.1 & \text{if } 1 - \frac{1}{9}(16t-11)^2 < 0.1 \\ 1 - \frac{1}{9}(16t-11)^2 & \text{if } 1 - \frac{1}{9}(16t-11)^2 > 0.1 \end{cases}
\end{aligned} \quad \text{(S.9)}$$

### Derivation of the reduced dimension repressilator

In order to reduce the dimensionality of the full repressilator system (Eqs. (6)-(9) in the main text), we followed the same quasi-steady state approximation (QSSA) introduced by Bennet et al. [2]. The initial set of equations is

$$\begin{aligned}
\dot{m}_i &= \alpha_i\pi_i - \delta_i m_i, && \text{(S.10)} \\
\dot{p}_i &= \sigma_i m_i - \gamma_i p_i + 2\kappa_i^- d_i - 2\kappa_i^+ p_i^2, && \text{(S.11)} \\
\dot{d}_i &= \kappa_i^+ p_i^2 - \kappa_i^- d_i - k_i^+ \pi_{i+1} d_i + k_i^-(1-\pi_{i+1}), && \text{(S.12)} \\
\dot{\pi}_i &= k_{i-1}^-(1-\pi_i) - k_{i-1}^+ d_{i-1}\pi_i. && \text{(S.13)}
\end{aligned}$$

The meaning of the parameters are detailed in the main text, and summarized in Fig. 4A. Following QSSA, we identified that the molecular reactions changing the state of each individual protein (monomer $p_i$, free dimer $d_i$, or bound dimer through the promoter occupancy $\pi_i$) can be assumed to be faster than the change of total protein in any form.

Therefore, $p_i$, $d_i$ and $\pi_i$ equilibrate faster than the other species, so we can assume that equations (S.11), (S.12) and (S.13) are at equilibrium. Setting $\dot{d}_i$ to zero, equation (S.12) gives us

$$\kappa_i^- d_i - \kappa_i^+ p_i^2 = k_i^-(1-\pi_{i+1}) - k_i^+ d_i \pi_{i+1}$$

The RHS of this equation is the RHS of equation (S.13) for $i \to i+1$. Since $\dot{\pi}_{i+1}$ is at equilibrium, we have

$$\begin{aligned}
\kappa_i^- d_i - \kappa_i^+ p_i^2 &= k_i^-(1-\pi_{i+1}) - k_i^+ d_i \\
&= \dot{\pi}_{i+1} \\
&= 0.
\end{aligned} \quad \text{(S.14)}$$

Therefore, after rearrangement,

$$d_i = \frac{\kappa_i^+}{\kappa_i^-} p_i^2. \quad \text{(S.15)}$$

Substituting this into equation (S.13), given that $\dot{\pi}_{i+1} = 0$ gives us

$$\begin{aligned} 0 &= \dot{\pi}_i \\ &= k^-_{i-1}(1-\pi_i) - k^+_{i-1} d_{i-1}\pi_i \\ &= k^-_{i-1}(1-\pi_i) - k^+_{i-1}\frac{\kappa^+_{i-1}}{\kappa^-_{i-1}} p^2_{i-1}\pi_i \end{aligned} \tag{S.16}$$

And so, after rearrangement

$$\pi_i = (1 + \frac{k_{+,i-1}}{k_{-,i-1}}\frac{\kappa_{+,i-1}}{\kappa_{-,i-1}} p^2_{i-1})^{-1}. \tag{S.17}$$

Setting $k_i \equiv \frac{k_{+,i}}{k_{-,i}}$, $\kappa_i \equiv \frac{\kappa_{+,i}}{\kappa_{-,i}}$, we have

$$d_i = \kappa_i p_i^2, \tag{S.18}$$

$$\pi_i = (1 + k_{i-1}\kappa_{i-1}p^2_{i-1})^{-1}. \tag{S.19}$$

For usual QSSA, at this point we would substitute these equilibrium values into equations (S.11) and (S.10), giving us a 6-species system. However, as Bennet points out [2], this treats $p_i$ as a solely 'slow' variable, even though it depends on dimerisation, a fast reaction. Instead, we define a new variable $n_i$, which is a truly 'slow' variable. The variable $n_i$ represents the total concentration of protein $i$ in any form, i.e. monomers, dimers, and those bound to regulatory sites. So,

$$\begin{aligned} n_i &= p_i + 2d_i + (1-\pi_{i+1}) \\ &= p_i + 2\kappa_i p_i^2 + k_i\kappa_i p_i^2(1 + k_i\kappa_i p_i^2)^{-1} \end{aligned} \tag{S.20}$$

Therefore,

$$\begin{aligned} \dot{n}_i &= \dot{p}_i \frac{\partial n_i}{\partial p_i} \\ &= \frac{\dot{p}_i}{\mu_i(p_i)} \qquad \text{with} \quad \mu_i(p_i) := 1/\frac{\partial n_i}{\partial p_i}. \end{aligned} \tag{S.21}$$

Then

$$\mu_i(x) = \left(\frac{\partial n_i}{\partial p_i}\right)^{-1} = \left(1 + 4\kappa_i p + \frac{2k_i\kappa_i p}{(1 + k_i\kappa_i p^2)^2}\right)^{-1}$$

In addition, from our definition of $n_i$ as the total concentration of protein $i$ in any form, we know that the only reactions that can change $n_i$ are protein degradation and translation. so:

$$\dot{n}_i = \sigma_i m_i - \gamma_i p_i$$

And so (substituting into Eq. S.21)

$$\frac{\dot{p}_i}{\mu_i(p_i)} = \sigma_i m_i - \gamma_i p_i$$

Together this gives us the reduced system:

$$\dot{p}_i = \mu_i(p_i)(\sigma_i m_i - \gamma_i p_i)$$

$$\dot{m}_i = \frac{\alpha_i}{1 + k_{i-1}\kappa_{i-1}p^2_{i-1}} - \delta_i m_i$$

Where

$$\mu_i(p) = \left(1 + 4\kappa_i p + \frac{2k_i\kappa_i p}{(1 + k_i\kappa_i p^2)^2}\right)^{-1}$$

We can rewrite these equations in terms of the non-dimensional variables $\tilde{p}_i = \frac{p_i}{p_i^*}$ and $\tilde{m}_i = \frac{m_i}{m_i^*}$, where $p_i^* = \frac{\sigma_i \alpha_i}{\delta_i \gamma_i}$ is the maximum value of $p_i$ in absence of repression (i.e. set $k_1^+ = k_2^+ = k_3^+ = 0$), and $m_i^* = \frac{\alpha_i}{\delta_i}$ is the maximum value of $m_i$ in the absence of repression. We also let $r_i \equiv \sqrt{\frac{\kappa_i}{k_i}}$, $s_i \equiv \sqrt{\kappa_i k_i}$, $\beta_i \equiv \frac{\sigma_i \alpha_i}{\delta_i \gamma_i}$.

Without loss of generality, we can drop the tildes. These changes give us the following system:

$$\begin{aligned} \dot{p_i} =& \mu_i \gamma_i (m_i - p_i) \\ \dot{m_i} =& \delta_i \left( \frac{1}{1 + (s_{i-1}\beta_{i-1}p_{i-1})^2} - m_i \right) \end{aligned} \tag{S.22}$$

Where

$$\mu_i = \left( 1 + 4\beta_i r_i s_i x_i + \frac{4\beta_i s_i^2 x_i}{(1 + (\beta_i s_i x_i)^2)^2} \right)^{-1}$$

and $r_i \equiv \sqrt{\frac{\kappa_i}{k_i}} = \sqrt{\frac{\kappa_{+,i}k_{-,i}}{\kappa_{-,i}k_{+,i}}}$, $s_i \equiv \sqrt{\kappa_i k_i} = \sqrt{\frac{\kappa_{+,i}k_{+,i}}{\kappa_{-,i}k_{-,i}}}$, $\beta_i \equiv \frac{\sigma_i \alpha_i}{\delta_i \gamma_i}$

### Parameter search details

| Symbol | Meaning | Value |
|---|---|---|
| $\alpha_i$ | Transcription rate | 20 |
| $\sigma_i$ | Translation rate | 10 |
| $\gamma_i$ | Protein degradation rate | 1 |
| $\delta_i$ | mRNA degradation rate | 3 |
| $\kappa_i^+$ | Protein dimerization rate | 10 |
| $\kappa_i^-$ | Dimer dissociation rate | 10 |
| $k_i^+$ | Binding of dimer to promoter rate | 10 |
| $k_i^-$ | Dissociation of dimer and promoter rate | 10 |

Table 1: Parameters for the repressilator used in the parameter search.

The parameter search was performed by generating 5000 sets of parameters perturbing a reference parameter system. The parameters for the reference system $r_i = r_i^{\text{ref}}$ can be found in Table 1. The perturbations from the parameters $r_i$ are expressed as $r_i = r_i^{\text{ref}} e^{\zeta_i}$, where $\zeta$ is a uniformly distributed random variable $\zeta_i \sim \text{Uniform}(-\log 4, \log 4)$. Each set of randomly picked parameters will correspond to a system (which can be thought of as the GRN for a simulated new species). We removed parameter sets that do not give oscillations, as these parameters perturb the system so much that the orbit is destroyed. The values of $r_1$, $r_2$ and $r_3$ respectively of the perturbed systems in Fig. 5F in the main text are shown in Table 2.

| Tempo | 4x | 2x | 1x | 0.5x | 0.25x |
|---|---|---|---|---|---|
| Orbit 1 | 0.25, 0.25, 0.25 | 0.50, 0.50, 0.50 | 1.00, 1.00, 1.00 | 2.00, 2.00, 2.00 | 4.00, 4.00, 4.00 |
| Orbit 2 | 0.25, 0.05, 0.25 | 0.26, 0.05, 0.26 | 0.50, 0.10, 0.50 | 1.00, 0.20, 1.00 | 2.00, 0.41, 2.03 |
| Orbit 3 | 0.31, 0.06, 0.06 | 0.50, 0.10, 0.10 | 1.00, 0.20, 0.20 | 2.00, 0.40, 0.40 | 4.00, 0.80, 0.80 |

Table 2: System parameters for Fig. 5F in the main text.

### Heterogeneity of rate $r_i$

The heterogeneity $\eta$ of the perturbed rates $r_i$ with respect to a reference system $r_i^{ref}$ was calculated by quantifying the deviation of the components $r_i^* = r_i / r_i^{ref}$ with respect to the homogeneous case $\vec{r^*} = (1, 1, 1)$. While there are different choices for this comparison, for the case of the repressilator, we found that the best correlation against prefactor heterogeneity was achieved through the angle $\theta$ between the homogeneous case $(1, 1, 1)$ and the vector $(\sqrt{r_1}, \sqrt{r_2}, \sqrt{r_3})$.

$$\eta = \tan^2 \theta = \frac{1}{\cos^2 \theta} - 1 = 3 \frac{r_1^* + r_2^* + r_3^*}{\left( \sqrt{r_1^*} + \sqrt{r_2^*} + \sqrt{r_3^*} \right)^2} - 1 \tag{S.23}$$

Hence, different deviations from the homogeneous case can be visualised as cones around the vector (1,1,1) (see Fig. S.2 A)

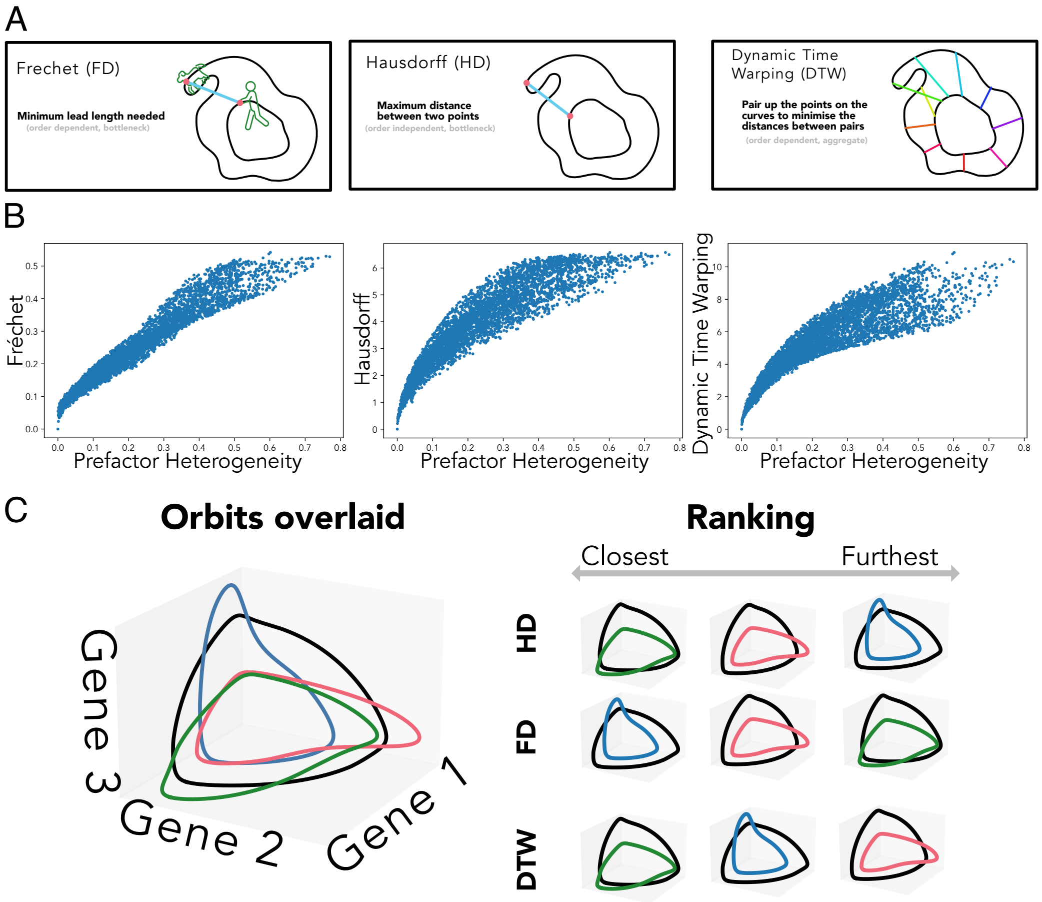


Figure S.1: A) Schematic comparing three different similarity measures. B) Comparisons of Fréchet, Hausdorff and dynamic time warping vs prefactor heterogeneity for the same perturbation set from Fig. 4C) Three different orbits in green, blue and pink are compared to a reference orbit (black curve). Different distance definitions rank the orbits in a different order.

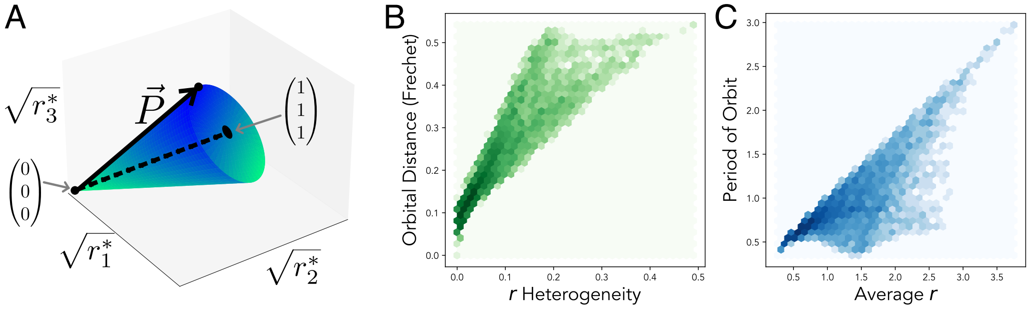


Figure S.2: A) A cone in the $r_i$ space is used to define $r$ heterogeneity. The angle $\theta$ measures the deviation of a point $\vec{P}$ from the homogenous direction $(1, 1, 1)$ (dashed line). B) $r$ heterogeneity vs the Fréchet distance for systems in the parameter search in Fig. 4 and Fig. 5. C) Average $r$ vs the period of orbit for systems in the parameter search in Fig. 4 and Fig. 5.
